# Spatial transmission dynamics of African Swine Fever in wild pigs: Sensitivity to epidemiological traits of different viral strains

**DOI:** 10.1101/2025.09.30.679520

**Authors:** Madison B. Berger, Ryan S. Miller, John M. Humphreys, Kim M. Pepin

## Abstract

African Swine Fever virus (ASFv) is a rapidly spreading animal disease with impacts on global economy and food security. Since the first known case of ASFv in Kenya in 1921, the virus has undergone significant evolution resulting in the emergence of 25 distinct genotypes. These genotypes have a wide variety of epidemiological traits, leading to uncertainty about how a genotype or variant of a genotype might spread in a population. Understanding the impacts of different epidemiological traits on the spread of ASFv is important for developing effective surveillance systems and control strategies. We developed a spatial epidemiological model of ASFv transmission in wild pigs, which contributes to ASFv persistence in countries where wild pig populations are widespread. We evaluated the impacts of a realistic set of epidemiological traits including incubation period, infectious period, disease-induced mortality rate, and length of immunity on spatial transmission dynamics under different ecological conditions for the host population (i.e. host density). We analyze effects of these conditions on outbreak metrics using boosted regression models with response variables: peak incidence, rate of spatial spread, seroprevalence and area invaded after one year, and probability that an outbreak will occur. We found that the infectious period of living individuals is the most important predictor at all pig densities for peak incidence and the area affected at one year, followed closely by the incubation period of the virus. We discuss considerations for surveillance and control strategies for different genotypes of ASFv.

## Introduction

African Swine Fever virus (ASFv) is an infectious disease that affects both domestic and wild pigs. Transmission routes include vehicles, contaminated feed or pork products, equipment, carcasses, or vectors such as ticks (Zhang, et al. 2023, Brown and Bevins 2018, Olesen, Lohse and Hansen, et al. 2018, Karalyan, et al. 2019, Bonnet, et al. 2020, Chenais, et al. 2018). An introduction of ASFv in countries that are currently free of the disease poses considerable impact on the pork industry and trade (Kashyap, Suter and McKee 2024). Strict regulations exist for imports and exports of all pork products to prevent the international exchange of contaminated products. Since 2005, 80 countries have reported ASFv cases with 9 of those being first ever occurrences (WOAH 2024). ASFv is predominantly found in Africa, central and southeast Asia and east Europe. The United States (U.S.) has remained disease-free largely due to its geographic distance from the ongoing epidemics and restrictions on importation of at-risk materials from affected countries. However, 40 years after being detected in the Western Hemisphere, the disease reappeared in both Dominican Republic and Haiti in 2021 (Schambow, et al. 2023, Jean-Pierre, Hagerman and Rich 2022, Gonzales, et al. 2021). This has led to an increase in surveillance and testing and questions about how ASFv might spread in the U.S. if an introduction were to occur.

Since the first observation of ASFv in the early 1900s, the virus has evolved due to genetic mutation and recombination (Sánchez-Cordón, et al. 2018, Chen, et al. 2024). There are currently 25 different genotypes of ASFv that vary in virulence and transmission with genotype II being the most lethal (Spinard, et al. 2023, Zhao, et al. 2023). All genotypes can be found circulating in Africa but only genotypes I and II have been detected outside of the continent (Hakizimana, et al. 2023). The strain that caused the outbreak in the Dominican Republic most closely resembles a milder form of genotype II from the Georgia 2007 outbreak, with 99.99% genetic similarity between the two isolates (Ramirez-Medina, et al. 2022). Mortality rate, incubation period, and infectiousness have shown to vary with these new strains (Sánchez-Cordón, et al. 2018, Chen, et al. 2024). Symptoms of ASFv can range from high fever, dyspnea, anorexia, nasal discharge, lesions, and cyanosis to death in acute cases (Sánchez-Vizcaíno, et al. 2015). The overall goal of this work is to explore how variation in mortality or other epidemiological parameters may impact spatial outbreak dynamics and conditions for surveillance.

Recent work synthesized the state of knowledge on epidemiological parameters for ASFv strains from experimental infection studies, including the duration of infectious and incubation periods, and transmission and recovery rates (Talbert, et al., 2025). This work provides a comprehensive overview of the variation in fundamental epidemiological traits among ASFv strains, irrespective of local host ecology. Their work focused on genotype II and found that most studies used the strain from the Georgia outbreak in 2007. However, outbreaks from as early as the 1950s have been characterized by reduced virulence and enhanced transmission (Olesen, Lohse and Boklund, et al. 2017, Ramirez-Medina, et al. 2022, Zani, et al. 2018, Wilkinson 1984).

In the current work, we use this representative view of variation in ASFv epidemiological traits to understand the possible range of spreading dynamics in the context of host ecology parameters that describe wild pig populations in the U.S. The overarching objective of this study is to examine effects of a realistic set of ASFv epidemiological parameters (as summarized in Talbert, et al., 2025) on spatial outbreak dynamics in wild pigs. Building on previous work (Pepin, et al. 2022), we tested the effects of host density, recovery and incubation periods, the infectious periods of both living wild pigs and carcasses, and mortality rates on predicting key outbreak traits. These traits include global seroprevalence at one year, peak incidence, the rate of spread across the landscape, the area affected at one year, and the probability of an outbreak occurring. This sensitivity analysis identifies the parameters with the most significant impact on disease outbreaks to provide insight for streamlining surveillance design for early detection and characterize the most important uncertainties for preparedness.

## Methods

We simulated ASFv transmission dynamics using an epidemiological metapopulation model previously described by Pepin, et al., 2022 which includes stochastic processes and host movement ecology. We evaluated the effects of realistic values of epidemiological parameters of ASFv on outbreak dynamics. Boosted regression trees were then used to determine which parameters had the greatest influence on response variables of interest.

### Simulation initialization

Six parameters were varied across our simulations. These are shown with corresponding values in Figure 1b. Because length of immunity is unknown, we investigated the effects of this parameter (ω) on epidemiological dynamics. Many of the more recently discovered genotypes have been shown to be less lethal than genotypes I and II. We investigate this process through parameter (□) - the proportion of pigs that recover from infection. Other parameters we varied, based on data from Talbert, et al., 2025, include average incubation period (σ), infectious period of both living individuals ( ) and carcasses ( ) as well as host density. To capture diverse epidemiological conditions, we considered all possible combinations of these parameters resulting in 7,200 distinct systems (3 host densities x 5 recovery proportions x 4 incubation periods x 5 carcass infectious periods x 4 living infectious periods x 6 recovery periods). For each parameter set, we ran 100 replicate simulations over a 72-week period (Gallardo, et al. 2017).

**Figure 1.**
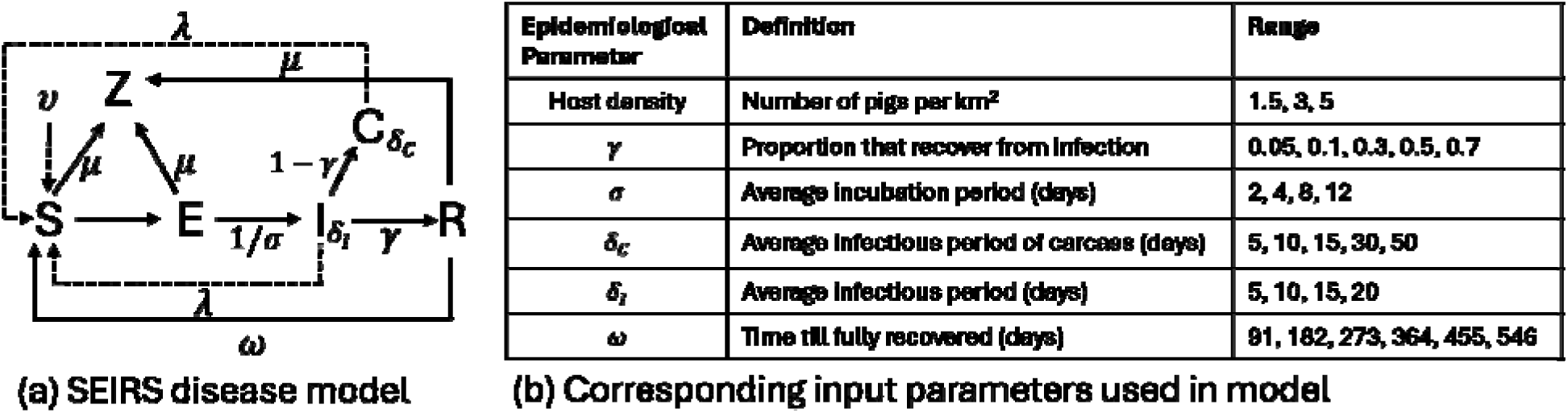
(a) Conceptual framework used to simulate the spread of African swine fever virus in wild pigs. States incorporated into this model include susceptible (S), exposed but not infectious (E), infectious and alive (I), non-infectious and recovered (R), non-infectious carcass within landscape (Z) and infectious carcass (C). (b) Discrete values for each epidemiological parameter used in simulations of disease spread.

To reflect key aspects of wild pig social structure in the model, we assumed a population organized into matriarchal groups (sounders), consistent with real world behavior where females and young males form groups and adult males are more solitary, though with overlapping home ranges (Maselli, et al. 2014). We implemented these dynamics using a spatially explicit metapopulation model, SQUEAL (Spatial QUantitative Evaluation for ASFv eLimination), in which each grid cell contained a sounder (see Pepin, et al., 2022 for details). A Poisson distribution based on average sounder size was used to determine the number of pigs per sounder and these were placed in grid cells that were randomly selected using a binomial distribution. The virus was introduced into one individual in the middle of an 80 km x 80 km homogeneous landscape and re-introduction of the virus was not allowed.

### SQUEAL model structure

The following disease states were used in this transmission model: susceptible (S), exposed (E), infectious (I), recovered (R), infectious carcass (C) and non-infectious carcass (Z) (Figure 1a). State transitions were implemented using a chain binomial model approach (Bailey 1957). An extension of Pepin, et al., (2022) incorporated in this study is the possibility for individuals to transition from recovered back to susceptible (*ω*). Two modes of transmission (λ) were incorporated into our model: 1) contact between two living individuals and 2) contact between a carcass on the landscape and a living individual. Carcass-based transmission has been shown to be a substantial mode of transmission, especially when accessing these carcasses on the landscape can be challenging (K. M. Pepin, A. J. Golnar, et al. 2020, Probst, et al. 2017, Cukor, et al. 2019, Rogoll, et al. 2024). We assumed individuals within the same grid cell had an equal probability of interacting with each other, *Φ* (Table S1). The probability that individuals will interact with carcasses is half of this same probability. This probability was based on the results of a spatially explicit epidemiological model fit to surveillance data that found that 53-66% of transmission events are carcass-based (K. M. Pepin, A. J. Golnar, et al. 2020). Between sounder transmission rates decayed with distance (Table S1).

We also allowed wild pig groups to shift home range centroids (moving to another grid cell). Such home range shifts were chosen from an empirical distribution of weekly home range shifts measured from wild pigs that were monitored with GPS collars in Florida, USA (Yang et al. 2021). In that population, weekly median location shifts for pigs were 163 m (Yang, Boughton, et al. 2021, Yang, Schlichting, et al. 2021). All cells containing pigs were selected and a home range shift was assigned to each grid cell. A home range shift occurred only if the movement distance was greater than 0.4 km which is equivalent to the resolution of the grid simulated on. Pigs would then be moved to a cell containing fewer pigs and within the distance assigned based on the gamma distribution of home range shifts (Table S1).

Births of susceptible (S) pigs occurred throughout the timeframe modelled with a limit of 10 per cell (V, Table S1) (Mayer and Brisbin 2009, Pepin, Wolfson, et al. 2019, Snow, Jarzyna and VerCauteren 2017). Natural death occurred in all disease states with a weekly binomial probability, μ (Table S1). Individuals that died either remained infectious (C) or not (Z) (Figure 1a). The disease-induced mortality rate was equivalent to 1-gamma (proportion that die from infection). Carcass persistence for both C and Z were modeled as a function of the infectious period, with the duration of persistence being influenced by the infectious period of the carcasses and transformed using a probabilistic exponential decay function.

### Analysis of simulation data

We selected five response variables that would provide a comprehensive view of ASFv outbreak progression, capturing key aspects such as disease spread speed, outbreak likelihood, scale and the extent of impact. Explicit response variables of interest included the area affected at 52 weeks, the global seroprevalence at 52 weeks, the median rate of spread in a week, peak incidence of the outbreak and the probability the outbreak will occur. A description of how each was calculated is shown in Table 1. The timeframe of 52 weeks was selected for several response variables because this would capture outbreak scenarios that were long lasting and not quick to progress through the population. All response variables except for outbreak probability were calculated only for outbreaks that occurred and do not account for stuttering or no further transmission past the first initial infection. The explicit criteria used to define an outbreak are shown in the last row of Table 1. The focus of this work is to understand disease spread therefore, removing results where outbreaks did not occur allowed for clearer interpretations of the modeling results.

**Table 1:** Description of each response variable and how it was calculated from simulations. Sero (+), refers to seropositive individuals that have been exposed or infected by ASFv and are producing antibodies in response. (*) Indicates that the response variable only included data from simulations that resulted in an outbreak.

| Response Variable | Description of Calculation |
| --- | --- |
| Area Affected at 52 Weeks*<br>(km<sup>2</sup>) | Area of a polygon surrounding all E, I, C and R individuals at 52 weeks. |
| Global Seroprevalence at 52 Weeks*<br>(sero (+) / Total Pop.) | The total number of R individuals (sero (+)) across the landscape divided by the total population of all living individuals (S, E, I, R) at 52 weeks. |
| Median Rate of Spread*<br>(km<sup>2</sup> / week) | Area of a polygon surrounding all E, I and C individuals at week $i$, subtracted by the area of a polygon surrounding all E, I and C individuals of the previous week ($i - 1$). The median rate of spread was selected from each outbreak. |
| Peak Incidence*<br>(cases) | The highest incidence that occurred in the outbreak. |
| Outbreak Probability | The proportion of simulation replicates classified as outbreaks, where an outbreak is defined as having a maximum number of incidences greater than 1 and a total number of incidences over the 72-week period greater than 10. |

We used gradient boosted regression tree (GBRT) models calculated using scikit-learn version 1.5.2 (Pedregosa, et al. 2011) to explore the complex interactions between our response variables and predictors. GBRT models use both regression tree algorithms and boosting which is used to combine the regression trees. Traditional regression methods construct a single model that provides the best results whereas boosted regression combines many simple trees adaptively which yields strong predictive models (Elith, Leathwick and Hastie 2008). Trees make decisions by splitting on individual predictors that best separate data at each stage. GBRT models do not explicitly calculate interaction terms rather, they identify predictor combinations that provide the best performance and least amount of error. The final feature importance score reflects the direct effect between a predictor and the response variable (Elith, Leathwick and Hastie 2008).

Model tuning involves modifying the hyperparameters used to construct simple trees. A *k*-fold cross validation technique was used to determine the best parameters for each regression model and to avoid overfitting (Yates, et al. 2022). This method involves splitting the dataset into 5 equal parts (or folds). For each iteration, the model is trained on 4 folds and tested on the remaining fold. This process is repeated until all folds have been used as the test set once. Data with the same predictor values were grouped together before being separated into folds to prevent the data with identical predictor values from showing up in the training data (Allgaier and Pryss 2024). This could lead to overfitting. A range of folds were tested (5-10) with ultimately 5 folds used based on the highest R^2^ value and computational time.

The optimal hyperparameters were selected based on the combination that yielded the highest R^2^ (Table S2). The number of trees corresponds to the number of boosting iterations used to build the final predictive model. We chose 1000 trees, as the large dataset allows for more iterations, enabling the model to capture more complex patterns. The max depth of each tree was set to 8 to prevent trees from becoming too deep, which helps reduce computational time and overfitting. A minimum of 6 samples was required at each node before it could be split into additional branches. We selected a low learning rate of 0.005, meaning that after each tree is constructed, the model updates the predictions by only 0.5% of the difference between the current prediction and the new tree’s prediction. A squared error loss function was used for all response variables, except for the probability of an outbreak, as it is commonly employed in regression tasks. For the outbreak probability, we used a log loss function due to the binary nature of the data, making it suitable for classification (De’ath 2007). Finally, when constructing the final regression model, the dataset was split into 80% for training and 20% for testing.

Finally, we used our regression trees to predict outbreak metrics for epidemiological parameters from several ASFv strains (Table 2). These strains are from outbreaks that occurred in Poland (POL/2015/Podlaskie/Lindholm), Estonia (EstoniaPol18_28298_O111), and the Dominican Republic (Zani, et al. 2018, Olesen, Lohse and Hansen, et al. 2018, Ramirez-Medina, et al. 2022). Notably, the Dominican Republic and Estonia strains are significantly less virulent than the Polish strain and have much longer infectious periods.

**Table 2:**
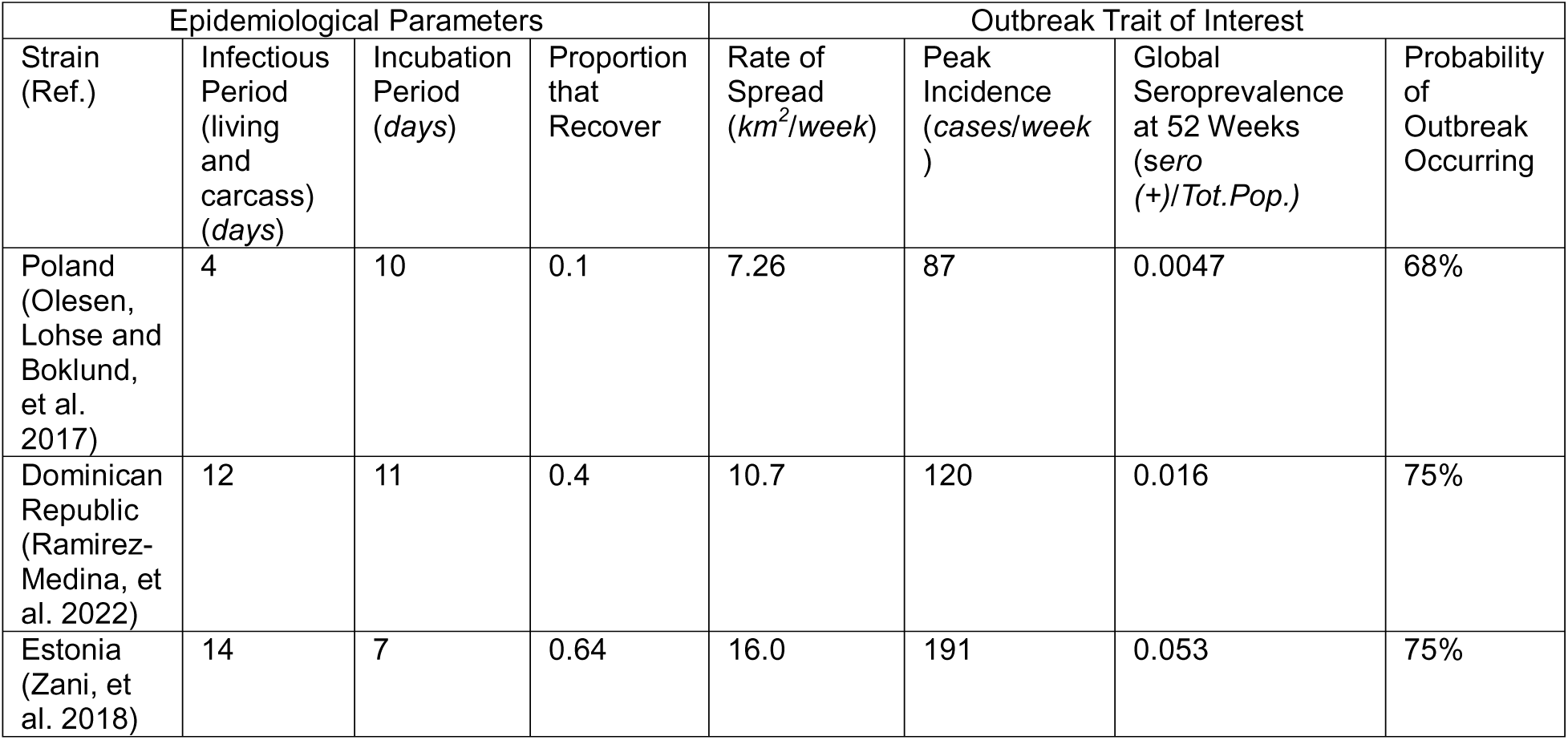
Using the gradient boosted regression models discussed, the epidemiological parameters from strains of ASFv found in the literature were used as predictors in the models using a medium host density. The recovery period was 546 days for all strains. Response variables are shown on the right side of the table.

## Results

### Infectious period of living

The strongest relationship between the predictors and incidence over time was infectious period of living animals (Figure 2, Figures S1-S4). As the infectious period increased, the rate of new cases increased at a much faster rate and this relationship was similar across medium and high host densities (Figure 2). Regression models built with respect to the rate of spread, peak incidence, area affected at 52 weeks, and outbreak probability all indicate the most important parameter driving these responses is the infectious period of the living (Figures 3-4, S5-S6, S8-S9). Specifically, the feature importance values for rate of spread were 0.78, 0.55, 0.39 for low, medium and high-density populations. At low host density, the relationship between rate of spread was linear across the discrete infectious periods we examined, and the range of rates became broader. This pattern was similar for medium to high densities, but the rate started to plateau at longer infectious periods. A similar plateau in higher host densities can also be seen in the area affected at 52 weeks regression model (Figure S8).

**Figure 2:**
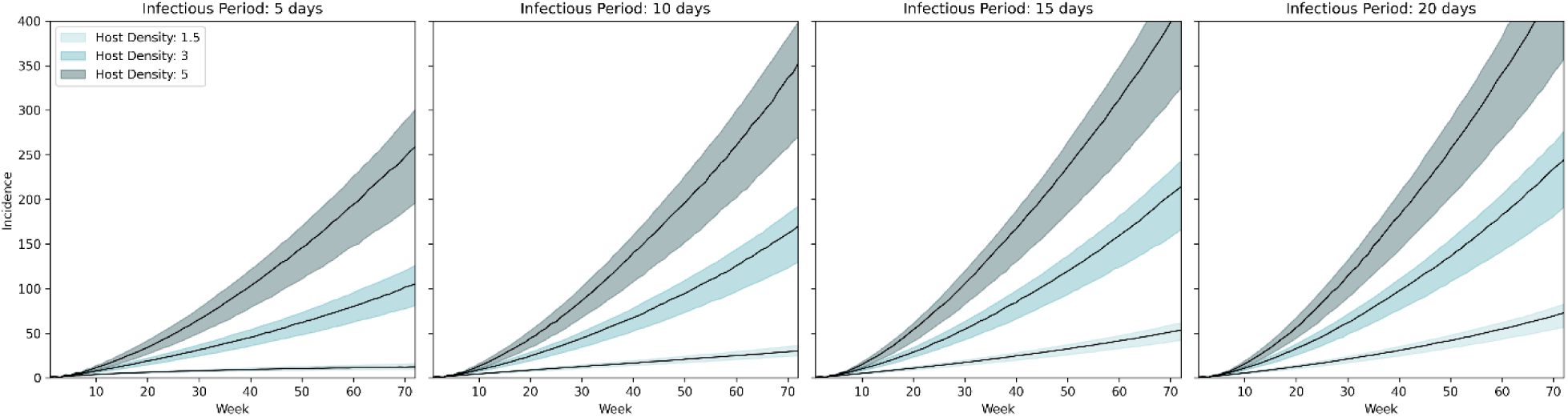
Incidence of new cases over time for each discrete infectious period of living individuals ( ). The black line represents the median number of new cases per week across all simulations, and the shaded region indicates the 50% confidence interval. Curves are grouped by host density.

**Figure 3:**
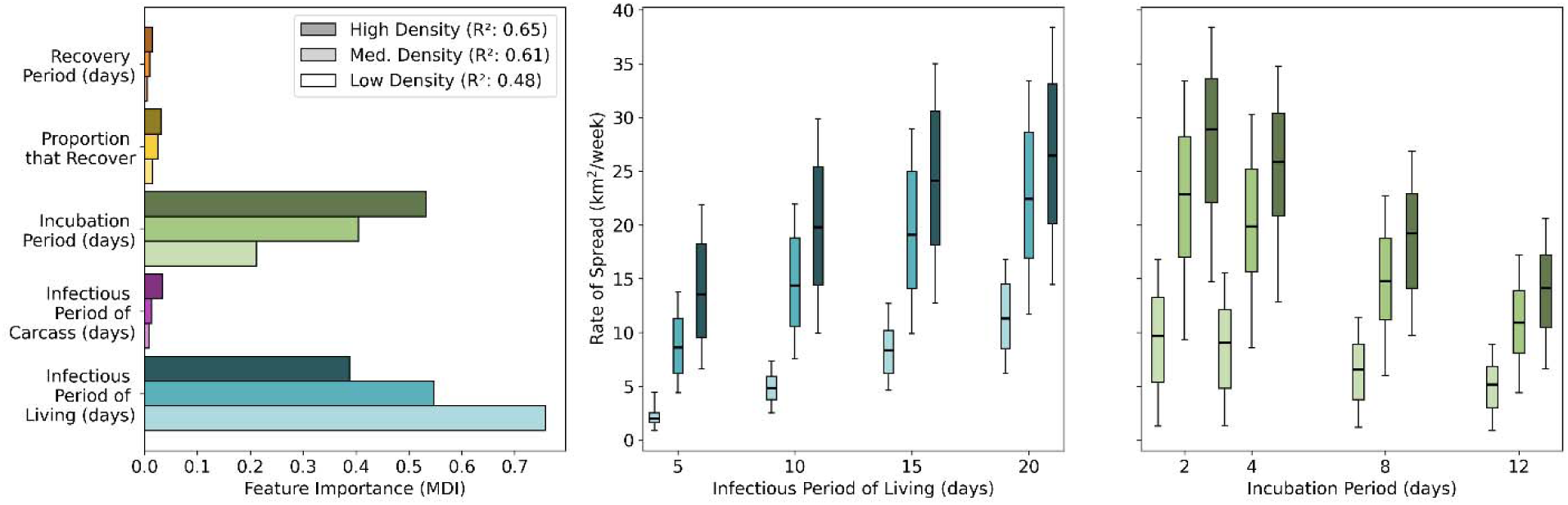
Results of a gradient boosted regression analysis in which the response variable was the rate of spread of individuals that were infectious (living and carcass) or exposed. The importance of each epidemiological trait is shown in the top left plot. The box and whisker plots indicate the relationship between each host density response and the epidemiological traits with high importance. The black lines on the boxes represent the median value. The whiskers indicate the maximum and minimum values. Low, medium and high-density populations are indicated by the light, medium and dark shaded bars respectively.

**Figure 4:**
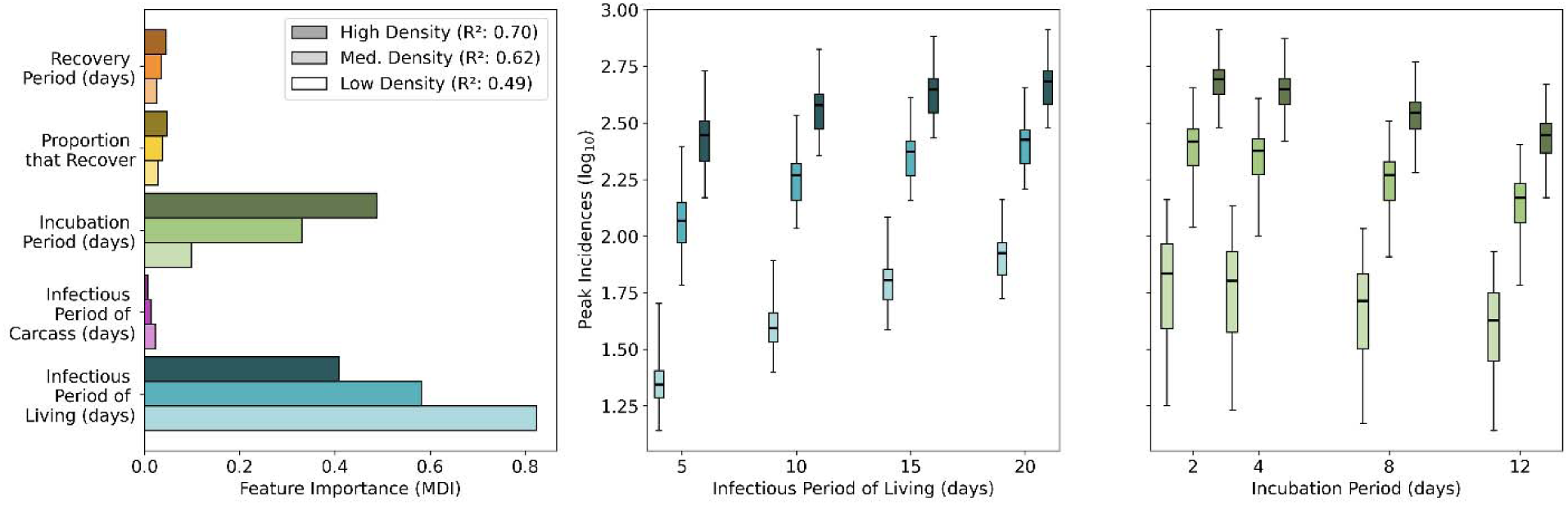
Results of a gradient boosted regression analysis. Response variable corresponds to the peak incidence of outbreaks on a log_10_ scale. The black line on the boxes represents the median value. The whiskers indicate the maximum and minimum values.

Similar trends were observed in the peak incidence regression model (Figure 4). The infectious period showed feature importances of 0.82, 0.58, and 0.41 for low, medium, and high-density populations. In low-density populations, the relationship between peak incidence and the infectious period was linear. However, at higher host densities, this relationship appeared to plateau, suggesting that moderate infectious periods led to rapid transmission, while longer infectious periods had diminishing effects on peak incidence (Figure 4). The duration of the infectious period also proved to be a significant factor in the outbreak probability regression model (Figure S9). While the infectious period was identified as one of the most important features, its importance was relatively close to that of other parameters in comparison to other regression models. This suggests that while the infectious period plays a key role, the probability of an outbreak is ultimately driven by the combined synergistic effects of all parameters.

### Infectious period of carcass

The infectious period of carcasses on the landscape is not a significant predictor in the outbreak response metrics explored here. This variable does show minor importance (<0.2 feature importance across all host densities) in influencing outbreak probability (Figure S9). Higher host densities are strongly associated with increased rates of spread, peak incidence, affected area, global seroprevalence, and overall outbreak probability.

### Incubation period

Incubation period was the second most significant predictor across all models, including those assessing the rate of spread, peak incidence, and the area impacted at 52 weeks (Figure 2, 3 and S8). An inverse relationship is observed between the length of the incubation period and the response variables. As the incubation period lengthens, the rate of spread, peak incidence, and total affected area tends to decrease.

### Proportion that recover

The recovery proportion plays a significant role in determining global seroprevalence at 52 weeks, with feature importance values of 0.71, 0.76, and 0.78 (Figure 5, S7). This predictor is consistently important at all host densities when compared to other variables in the regression models. As the proportion of individuals who recover increases, seroprevalence shows a slight increase. However, it is important to note that our model does not account for herd immunity because little is known about ASFv immunity after recovery. Therefore, Figure 5 does not imply that recovery directly reduces transmission or provides population-level immunity. Because the model does not consider herd immunity, the increase in global seroprevalence reflects the cumulative recovery of individuals, without reducing transmission or affecting broader outbreak dynamics. Recovery proportion minorly impacts outbreak probability (Figure S9) with a decrease in outbreak likelihood as the recovery proportion increases. This could be caused by a saturation of individuals recovering and thus, no individuals available to become infected.

**Figure 5:**
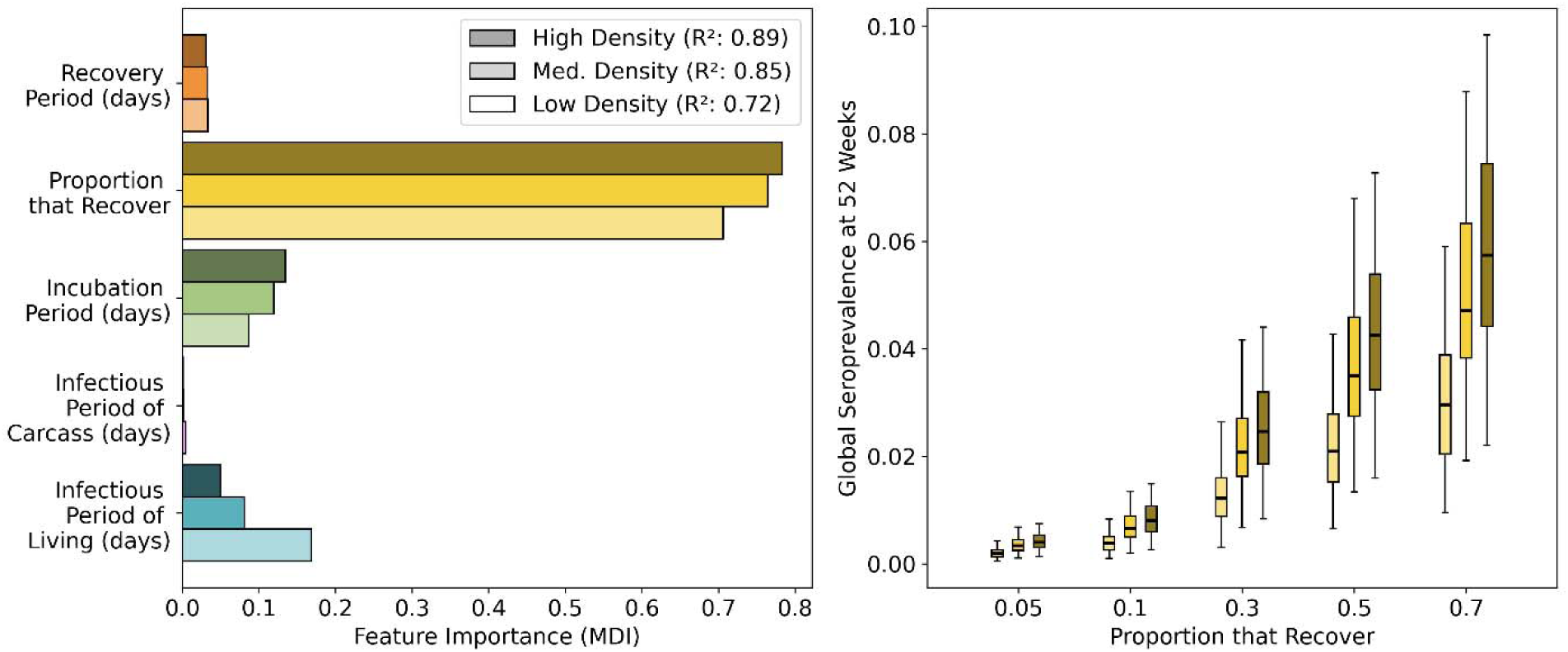
Results of a gradient boosted regression analysis. Response variable is the global seroprevalence at one year. The black line on the boxes represents the median value. The whiskers indicate the maximum and minimum values.

### Recovery Period

The recovery period is not a major predictor for any of the response variables in our regression models (<0.1 feature importance). This is largely because our model does not incorporate herd immunity. Without accounting for immunity, the recovery period plays more of a secondary role in the disease dynamics. While it affects the duration of recovery, it does not directly influence transmission dynamics or the spread of the disease in the population. As a result, the recovery period does not significantly impact key outbreak parameters such as the rate of spread, peak incidence, or affected area, as these are driven by other factors such as host density and the infectious period.

### Application to real-world strains

Using parameters from real-world strains evaluated in experimentally (i.e., irrespective of local host ecology), when combined with U.S.-based wild pig ecology we found that longer infectious periods substantially increase the rate of spread, peak incidence, and outbreak probability compared to the Polish strain (Figures 3, 4, S9 and Table 2). Additionally, the higher recovery proportion for both the Dominican Republic and Estonia strains contributes to a greater global seroprevalence, as predicted by our model (Figure 5, Table 2). In conclusion, the less virulent strains such as Estonia are associated with faster rates of spread, higher peak rates of new cases, and more widespread seroprevalence across the population at one year compared to the more virulent Polish strain.

## Discussion

The sensitivity analysis conducted in this study provides insights into viral parameters that may influence the spread of ASFv among wild pigs. Our results highlight that the infectious period, incubation period, and recovery proportion have the largest impact on determining outbreak dynamics, given our model assumptions and the response variables considered. These findings underscore the importance of thoroughly understanding the epidemiological characteristics of ASFv during an outbreak. In addition to these viral traits, host density in the affected area is a crucial component to consider, as higher densities can facilitate the rapid transmission of the virus, making it a critical factor to consider when modeling outbreak scenarios (K. M. Pepin, A. J. Golnar, et al. 2020, Podgórski, et al. 2020). The findings presented in this study support this relationship, showing that areas with higher wild pig densities experienced more rapid viral spread. However, to further refine our models and enhance predictive accuracy, it is essential to gather more comprehensive data on the full spectrum of ASFv strains. Expanding these data will improve our ability to accurately predict and control ASFv spread. With these improvements, our model can better inform surveillance and control efforts during emergency response situations (Rahman, et al. 2019, Galvis, et al. 2022).

The observed linear relationship between host density and infectious period of living individuals indicates that as population density increases and individuals remain infectious for longer periods, there are more opportunities for transmission (de Carvalho, et al. 2013). This amplification of transmission risk can drive outbreaks more quickly, emphasizing the need for swift characterization of these epidemiological traits following the virus’s introduction (Schulz, et al. 2019, O’Neill, et al. 2020). However, as host density and the duration of infectious periods rise, the affected area and rate of spread begin to plateau (de Carvalho, et al. 2013, Halasa, et al. 2016, Lentz, et al. 2023). This suggests that beyond a certain point, longer infectious periods may no longer significantly affect these outcomes, likely because of limitations due to host factors such as density, movement, or contact rates.

ASFv-specific modeling quantifying the impact of viral incubation period on outbreak dynamics is limited. General epidemic simulations demonstrate that long incubation periods tend to produce slower, flatter incidence curves with lower peaks and delayed timing (Kahn, et al. 2020, Arzt, et al. 2019). Our simulation results similarly suggest that strains with prolonged incubation periods could reduce the transmission rate leading to a lower peak incidence and less widespread impact. Longer incubation periods would provide more time for emergency management which would in turn, limit the overall spatial scope of the outbreak.

Although recovery time and infectiousness of carcasses showed low sensitivity in our analysis suggesting they have less influence on epidemic dynamics, they still warrant consideration in the broader context of disease management strategies. Effective carcass disposal and understanding of recovery timelines are crucial for mitigating indirect transmission pathways and ensuring appropriate control measures (Arzumanyan, et al. 2021, Eblé, et al. 2019). Our model assumes that the probability of contact with carcasses is half that of contact with living animals, which greatly affects the infectious period of carcasses. An increase in the contact rate could elevate the importance of this parameter. Previous studies have indicated that approximately half of transmission events are carcass-based, with this type of transmission increasing as host density decreases (Pepin, Golnar and Podgórski 2021). Therefore, our assumption of 50% carcass-based transmission might oversimplify this dynamic. This form of transmission could be heavily influenced by environmental conditions, rate of decay, or the social behavior of pigs. Further observations of living pigs interacting with carcasses are essential to our interpretation of this model. Additionally, GBRT models can explore how combinations of two predictors might be related in an additive manner. Multiplicative interactions are not explicitly captured as well as non-linear relationships between predictors (Natekin and Knoll 2013). There could be non-linear interactions occurring between the infectious period of carcasses or recovery time with other predictors explored in this work that the model was unable to capture.

Previous work has shown that pig populations with higher movement rates can yield similar qualitative results to populations with lower movement rates, as used in this study (Pepin, et al. 2022). However, the quantitative outcomes differ, with higher movement populations leading to more individuals being infected. We anticipate that incorporating higher movement dynamics would result in similar trends in the sensitivity analysis, but with increases in response variables, such as spread rate and affected geographic area. These results reinforce the need to consider population movement patterns when modeling ASFv spread, as omitting movement could lead to underestimating outbreak intensity and extent.

### Implications for surveillance design

As the proportion of individuals that recover increases and recovery time lengthens, serological testing becomes an increasingly valuable tool for ASFv detection. In contrast, when strains are highly lethal, carcass surveillance remains a critical component of early detection, as deaths occur more rapidly (Guinat, et al. 2016, Allepuz, et al. 2022, Gervasi, et al. 2020). Strains with longer infectious periods could delay the detectability of the virus via PCR, highlighting the importance of increasing sampling frequency. Additionally, strains with prolonged infectious periods tend to spread more rapidly and affect larger areas, suggesting that expanding surveillance across a broader spatial range could improve sampling effectiveness. This work also showed that peak incidence is higher for strains with extended infectious periods, implying that the total number of samples may be less important than focusing on the region sampled and the frequency of sampling (Schulz, et al. 2019). Many of these surveillance strategies assume that managers are familiar with epidemiological parameters of the strain causing the outbreak. Therefore, it is essential for researchers to understand both the likely points of introduction and the origin region, as this knowledge will help guide decisions regarding strain characteristics and surveillance approaches.

### Limitations of model design

In this study, we examined a variety of relevant response variables that offer valuable insights into the dynamics of ASFv outbreaks. However, several additional metrics such as the basic reproduction number (R), outbreak duration, and changes in population size could further enhance our understanding of disease progression. For example, longer infectious periods may increase both R_0_ and outbreak duration by saturating the landscape with infectious individuals allowing more opportunities for transmission and longer outbreaks.

Outbreak dynamics are inherently time-dependent and complex. In this model, disease spread was simulated over a 72-week period, which restricts our ability to capture the full progression of outbreaks, particularly those that are either still growing or declining. Extending the simulation over a longer time frame or until the outbreak fades out (e.g., two years) would allow us to explore how these epidemiological traits influence outbreaks across a more extended period.

Another factor not currently captured in our simulations is the effect of viral re-introductions. Re-introductions have shown to play a key role in the prolonged persistence of several strains of ASFv such as the prominent strain in Poland (K. M. Pepin, A. J. Golnar, et al. 2020). The current implementation does not include additional introductions of the virus past the first week, which may underestimate the persistence of certain outbreak scenarios.

Our transmission model assumes a homogeneous landscape and does not incorporate habitat-driven movement, rather, sounders move based on the absence of other pigs in neighboring areas and the probability of a weekly location shift. While this approach allows us to isolate the influence of viral epidemiological traits on outbreak dynamics, it does not include key ecological drivers of movement such as habitat preference and heterogeneity, and how wild pigs move in different habitat structures. If landscape features were to restrict sounders from moving to other regions, this could result in clustering within a given area, increasing the likelihood of local transmission, or epidemic fadeout when large patches of poor quality habitat are present. For instance, if an infectious pig dies in a densely populated or high-movement area, the probability of carcass transmission could be elevated. Habitat-driven movement could be further explored by expanding our movement function to include habitat selection and running the simulations on heterogeneous landscapes. We are currently pursuing this work to understand how surveillance design and response could be optimized in different ecological contexts across the U.S.

Finally, herd immunity remains poorly understood in the context of ASFv, as noted in the existing literature (Sauter-Louis, et al. 2021, Orosco 2023). In our model, individuals can only transition from recovered to susceptible or deceased, which results in a continuous increase in the susceptible population across the landscape, with a lag while individuals are temporarily immune. This period of immunity can be long based on the recovery period parameter that was varied (13-78 weeks). However, births occur every week allowing for a steady number of susceptible individuals. The long recovery period facilitates the potential for persistence. While immunity also could be achieved through vaccination, this was not considered in our model due to the current absence of an approved vaccine for ASFv (Ntakisyisumba, Tanveer and Won 2025). As research advances and more data become available on the effectiveness of vaccines or length of immunity from natural infection, these aspects will be further understood.

## Conclusion

This analysis provides a deeper understanding of how epidemiological traits contribute to ASFv outbreak dynamics. The infectious period of living individuals, incubation period, and recovery proportion are key drivers in the area affected at 52 weeks, global seroprevalence at 52 weeks, the rate of geographic spread, peak incidence, and the probability of an outbreak. Our findings suggest that while less virulent ASFv strains might result in less severe individual infections, they may contribute to larger outbreaks overall due to their prolonged infectious periods and higher recovery proportion that keep individuals contributing to new births of susceptible individuals in the population. This finding could have significant implications for surveillance design and biosecurity policies, as less virulent strains may spread more widely and rapidly, despite their lower individual-level severity.

## Acknowledgements

MBB is a fellow within the Agricultural Research Service (ARS) Research Participation Program which is coordinated by the Oak Ridge Institute for Science and Education (ORISE). This project was funded in part by the U.S. Department of Agriculture, Agricultural Research Service-CRIS project 3022-32000-064-000-D, Intervention Strategies to Respond, Control, and Eradicate Foot-and-Mouth Disease Virus (FMDV). This research used resources provided by the SCINet project of the USDA Agricultural Research Service, ARS project numbers 0201-88888-003-000D and 0201-88888-002-000D. KMP was supported by the United States Department of Agriculture, Animal and Plant Health Inspection Service, Wildlife Services. Any use of trade, firm, or product names is for descriptive purposes only and does not imply endorsement by the U.S. Government. The findings and conclusions in this article are those of the authors and do not necessarily represent the views of the US government. The authors would like to thank Raoul Boughton for contributing the foundational dataset allowing this work to be possible and Kayleigh Chalkowski for insightful conversation.

## Data Availability

Code and inputs used to generate the model output data presented in this manuscript have been archived on figshare (DOI 10.6084/m9.figshare.28811111). Code for all analyses discussed is provided at https://github.com/geoepi/ASF-GBRTs.

## Notes

### Competing Interest Statement

The authors have declared no competing interest.

