## Supplementary Information for "Spatial transmission dynamics of African Swine Fever in wild pigs: Sensitivity to epidemiological traits of different viral strains"

| **Epidemiological Parameter** | **Value** |
| --- | --- |
| $\upsilon$, Weekly birth rate | 0.0192 week^-1^ |
| $\mu$, Weekly natural death rate | 0.0064 week^-1^ |
| Weekly location shift distance | ~gamma(0.355, 0.751) km |
| Initial mean family group size | 2, 4 or 6 pigs per grid cell based on density (1.5, 3 or 5) |
| $\lambda$, Transmission rate (direct and carcass-based) | 0.74 |
| *Φ,* Scaling parameter for transmission probability with contact | 0.9, 0.4, 0.2 for 1.5, 3 and 5 host density |
| Carcass-based scaling parameter (each density) | *Φ* x 0.5 |
| Direct contact rate among grid cells | *α*=-0.80, *β*=-1.91 |
| Carcass-based contact rate among grid cells | *α*=2.32, *β*=-6.37 |

**Table S1**: Summary of fixed parameters used in simulation model. Parameters are based on Table 1 from Pepin, et al., 2022.

| **Response Variable** | **Number of Trees** | **Max Depth** | **Minimum Samples Split** | **Learning Rate** |
| --- | --- | --- | --- | --- |
| Area Affected at 52 Weeks | 1000 | 8 | 6 | 0.005 |
| Global Seroprevalence at 52 Weeks | 1000 | 8 | 6 | 0.005 |
| Rate of Spread | 1000 | 8 | 6 | 0.005 |
| Peak Incidences | 1000 | 8 | 6 | 0.005 |
| Outbreak Probability | 1000 | 8 | 6 | 0.005 |

**Table S2**: Hyperparameters used to build regression models.

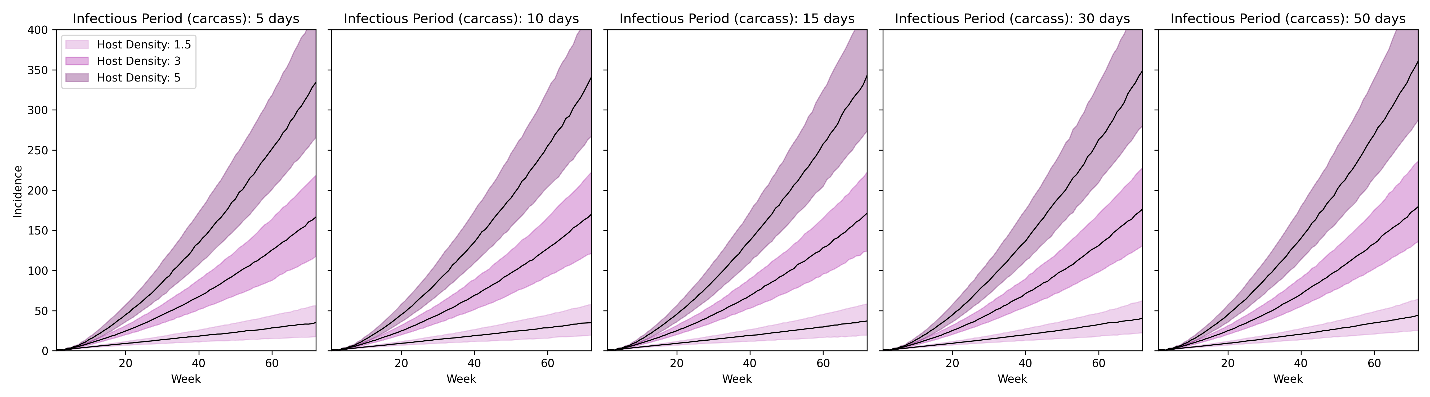
**Figure S1**: Incidence of new cases over time for each discrete carcass infectious period ($\delta_{c}$). The black line represents the median number of new cases per week across all simulations, and the shaded region indicates the 50% confidence interval. Curves are grouped by host density.

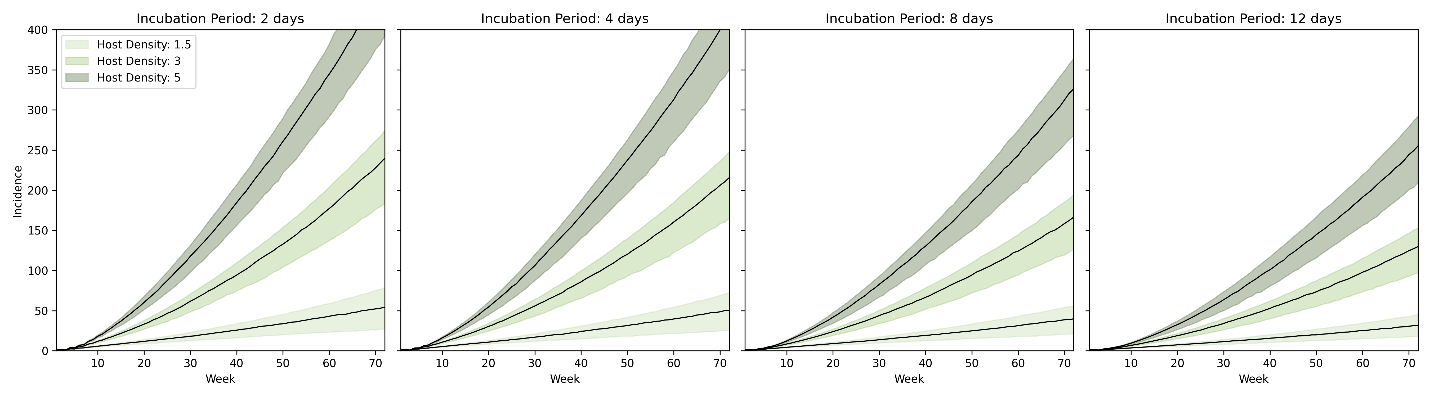
**Figure S2**: Incidence of new cases over time for each discrete incubation period (σ). The black line represents the median number of new cases per week across all simulations, and the shaded region indicates the 50% confidence interval. Curves are grouped by host density.

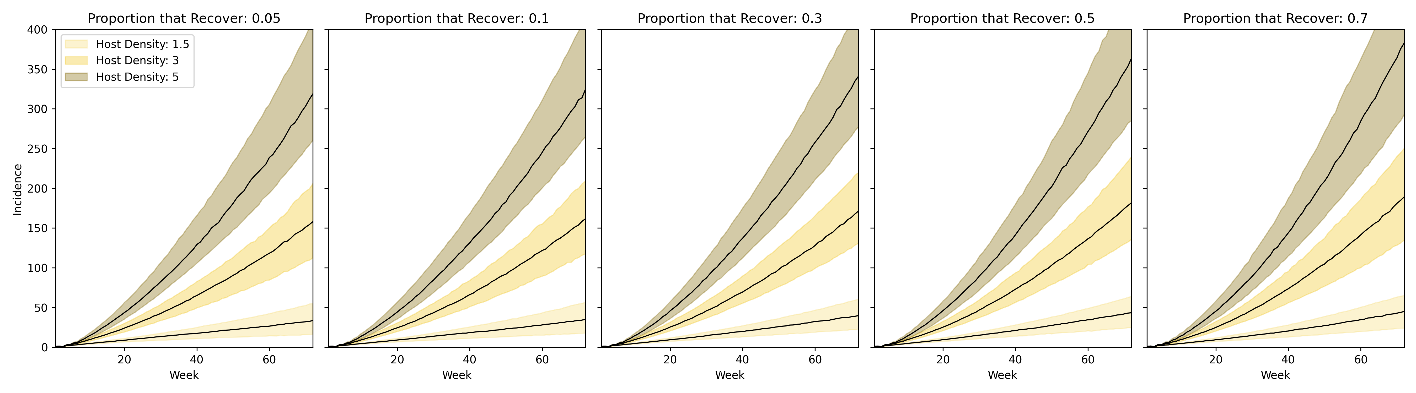
**Figure S3**: Incidence of new cases over time for each discrete proportion of individuals that recover (ɣ). The black line represents the median number of new cases per week across all simulations, and the shaded region indicates the 50% confidence interval. Curves are grouped by host density.

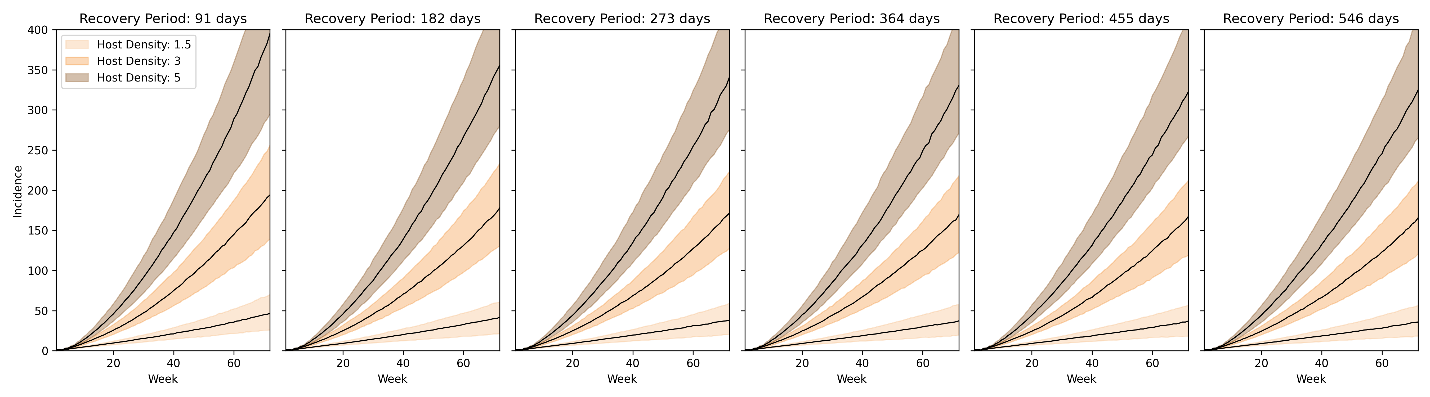
**Figure S4**: Incidence of new cases over time for each discrete recovery period (ω). The black line represents the median number of new cases per week across all simulations, and the shaded region indicates the 50% confidence interval. Curves are grouped by host density.

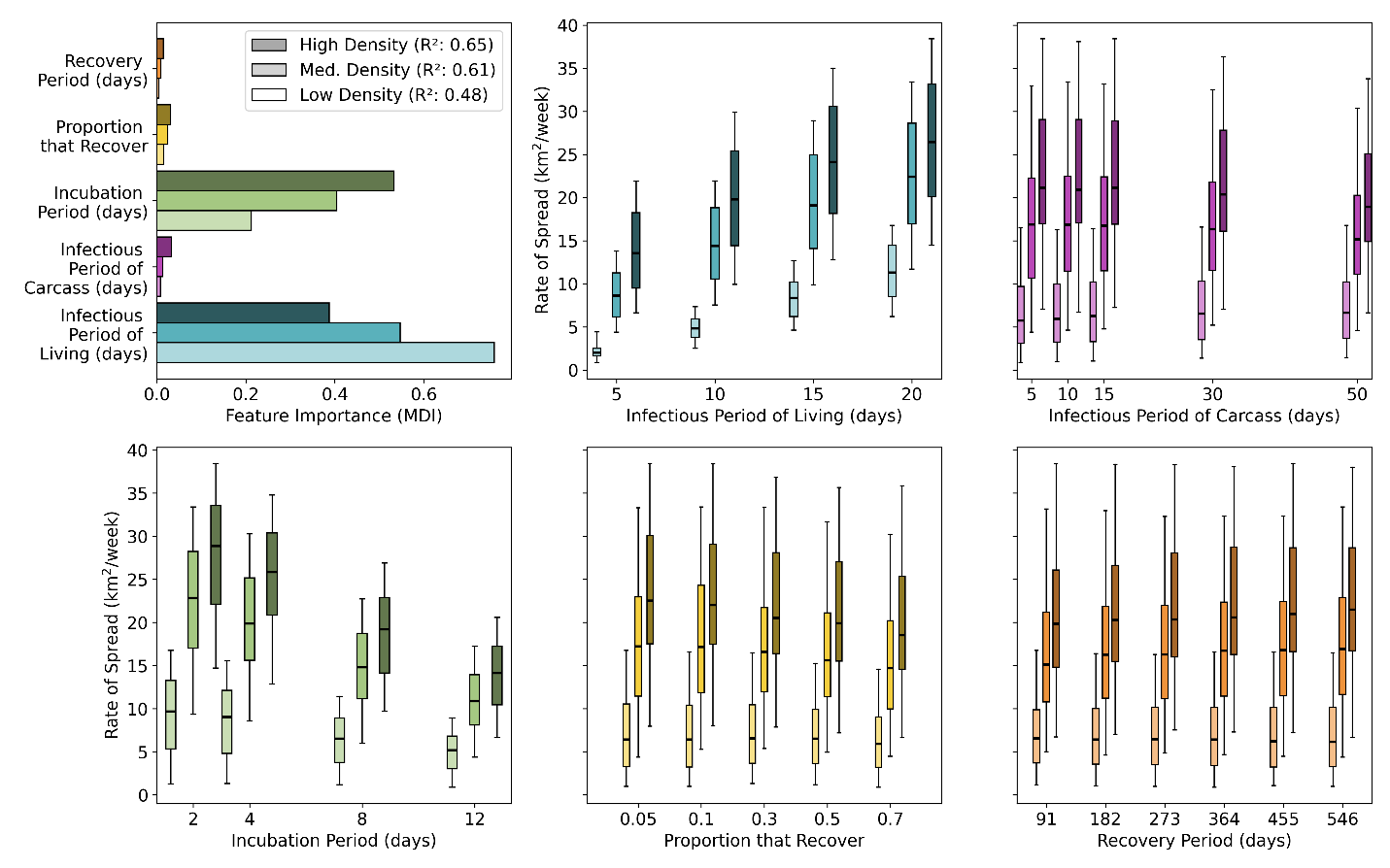

**Figure S5:** Results of a gradient boosted regression analysis in which the response variable was the rate of spread of individuals that were infectious (living and carcass) or exposed. The importance of each epidemiological trait is shown in the top left plot. The box and whisker plots indicate the relationship between each host density response and all epidemiological traits. The black line on the boxes represents the median value. The whiskers indicate the maximum and minimum values. Low, medium and high-density populations are indicated by the light, medium and dark shaded bars respectively.

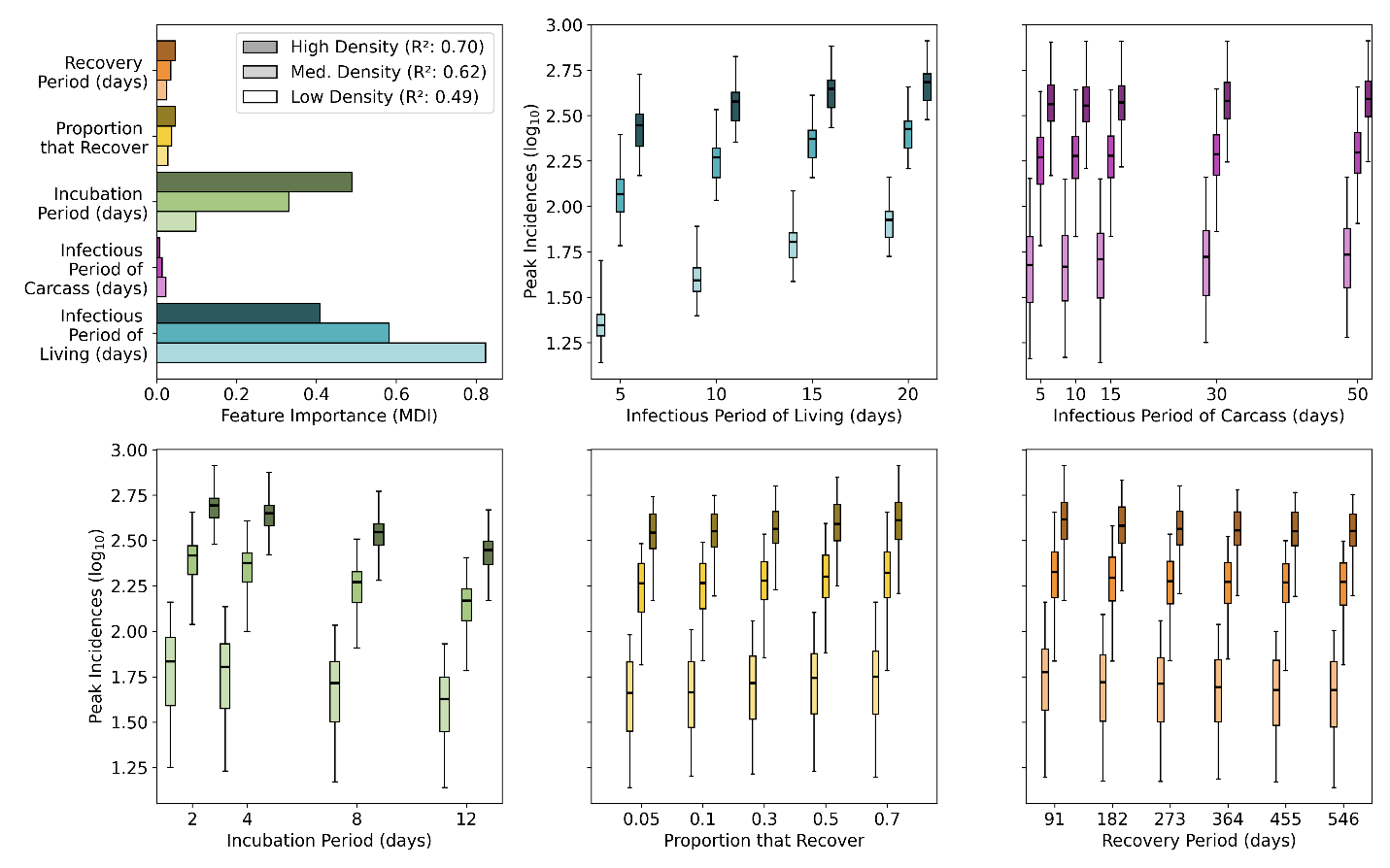

**Figure S6:** Results of a gradient boosted regression analysis in which the response variable was the peak incidence of outbreaks on a log_10_ scale. The importance of each epidemiological trait is shown in the top left plot. The box and whisker plots indicate the relationship between each host density response and all epidemiological traits. The black line on the boxes represents the median value. The whiskers indicate the maximum and minimum values. Low, medium and high-density populations are indicated by the light, medium and dark shaded bars respectively.

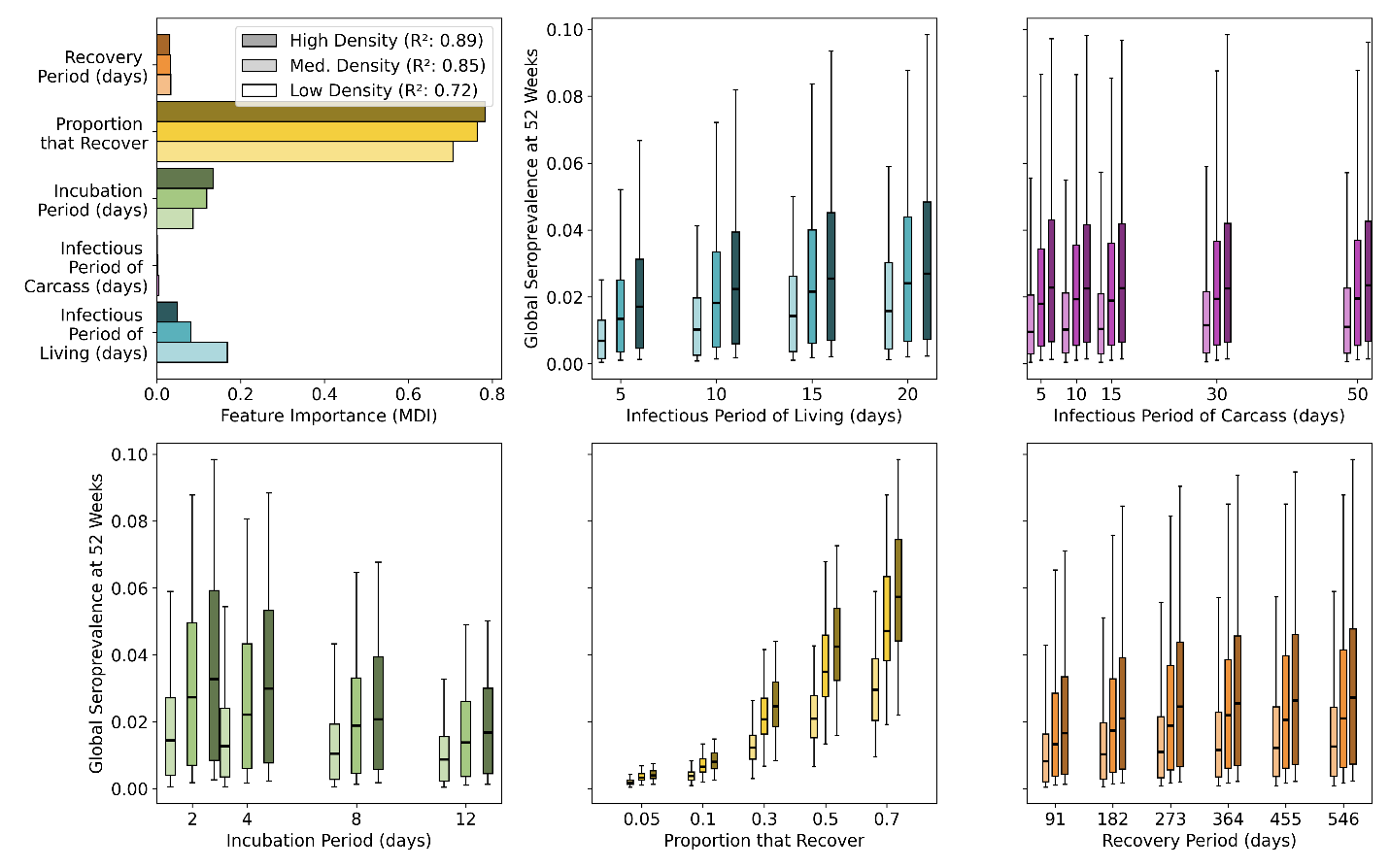

**Figure S7:** Results of a gradient boosted regression analysis in which the response variable is the global seroprevalence at one year. The importance of each epidemiological trait is shown in the top left plot. The box and whisker plots indicate the relationship between each host density response and all epidemiological traits. The black line on the boxes represents the median value. The whiskers indicate the maximum and minimum values. Low, medium and high-density populations are indicated by the light, medium and dark shaded bars respectively.

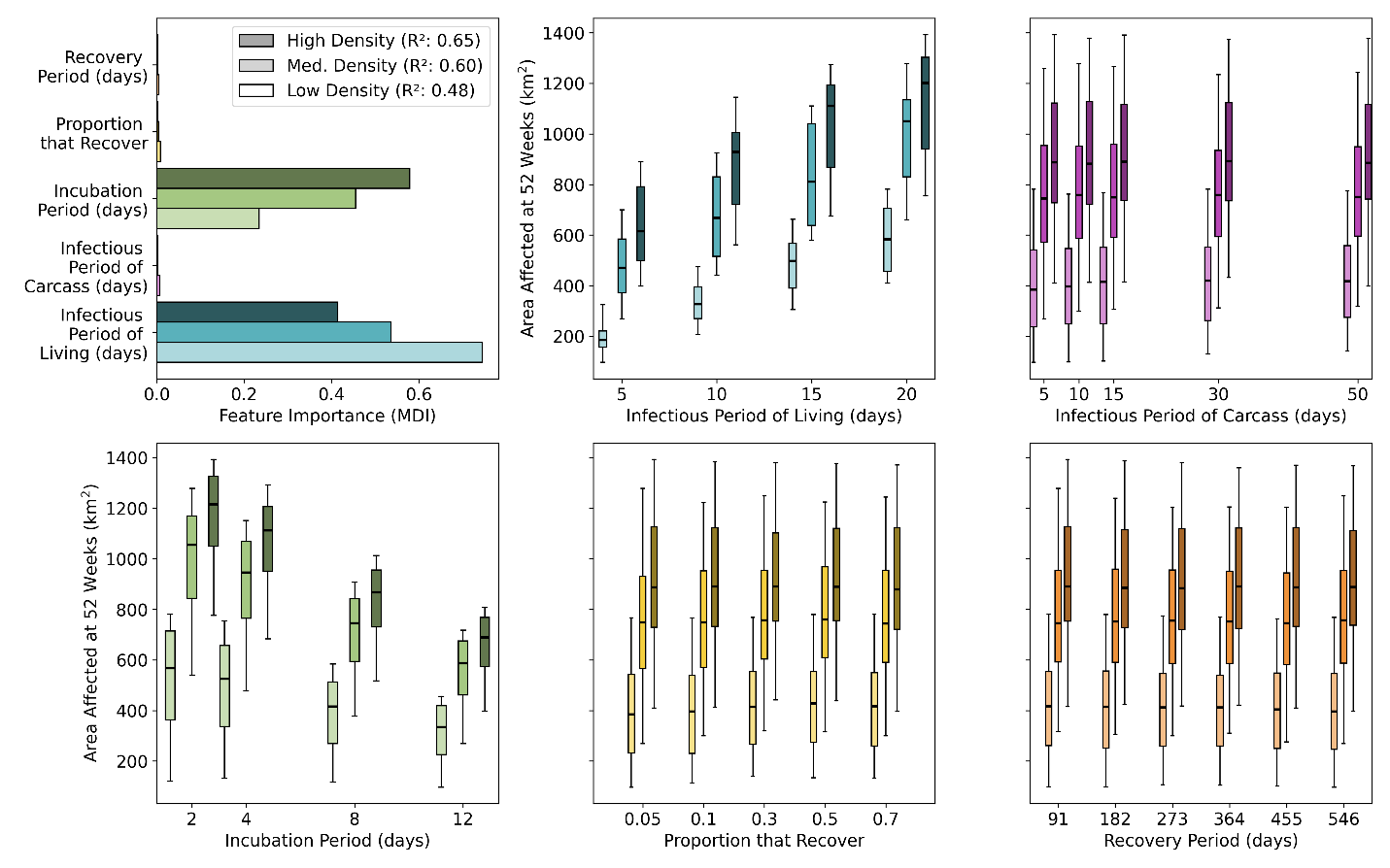
**Figure S8**: Results of a gradient boosted regression analysis in which the response variable is the area affected at one year. The importance of each epidemiological trait is shown in the top left plot. The box and whisker plots indicate the relationship between each host density response and all epidemiological traits. The black line on the boxes represents the median value. The whiskers indicate the maximum and minimum values. Low, medium and high-density populations are indicated by the light, medium and dark shaded bars respectively.

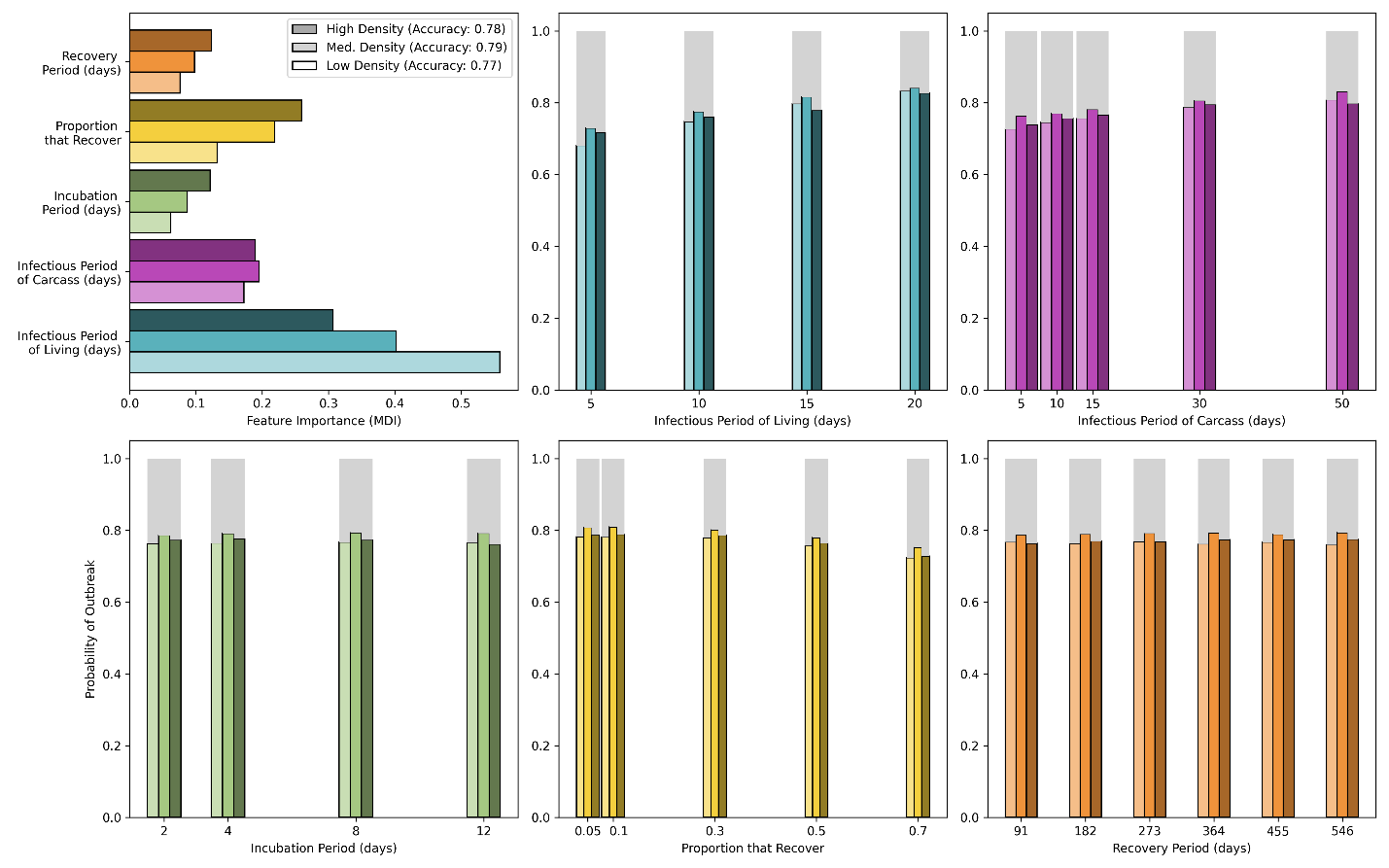

**Figure S9**: Results of a gradient boosted regression analysis in which the response variable is probability of an outbreak occurring. The importance of each epidemiological trait is shown in the top left plot. The univariate plots indicate the probability an outbreak will occur with respect to each discrete epidemiological trait value. The gray section indicates the probability that an outbreak will not occur.
